# Characterization and expression QTL analysis of *TaABI4*, a pre-harvest sprouting related gene in wheat

**DOI:** 10.1101/2020.09.18.303065

**Authors:** Chunsheng Xiao, Yujiao Liu, Wenshuai Chen, Jian Yang, Mengping Cheng, Calum Watt, Jingye Cheng, Zhenzhong Wang, Zhi Tan, MaoLian Li, Jirui Wang

## Abstract

Pre-harvest sprouting (PHS) induced by the decline of seed dormancy causes a severe reduction in crop yield and flour quality. In this study, we isolated and characterized *TaABI4,* an ABA-responsive transcription factor that participates in regulating seed germination in wheat. Sequence analysis revealed that *TaABI4* has three homologues, located on chromosomes 1A/1B/1D. TaABI4 contains a conserved AP2 domain, and AP2-associated, LRP, and potential PEST motifs. Putative *cis*-acting regulatory elements (CE1-like box, W-box, ABRE elements, and RY-elements) were identified in the *TaABI4* promoter region that showed high conservation in 17 wheat cultivars and wheat-related species. Expression profiling of *TaABI4* indicated that it is a seed-specific gene accumulating during the middle stages of seed development. Transcript accumulation of *TaABI4* in wheat cultivar Chuanmai 32 (CM32, PHS susceptible) was 5.07-fold and 1.39-fold higher than that in synthetic hexaploidy wheat SHW-L1 (PHS resistant) at 15DPA and 20DPA, respectively. Six expression quantitative trait loci (eQTL) of *TaABI4* on chromosome 2A, 2D, 3B, and 4A were characterized based on the accumulated transcripts of *TaABI4* in SHW-L1 and CM32 derived recombinant inbred lines. These QTLs explained from 10.7% to 46.1% of the trait variation with 4.53~10.59 of LOD scores, which contain genes that may affect the expression of *TaABI4*.

## Introduction

Pre-harvest sprouting (PHS) is the germination of mature grains in the spike when there is excessive moisture before harvest. PHS has become a recurring worldwide problem since it causes a severe reduction in crop yield and flour quality due to starch and protein degradation (Olaerts et al., 2016). Seed dormancy accounts for up to 60% of the variation in PHS tolerance, and PHS in wheat is mainly caused by the lack of adequate seed dormancy (DePauw and McCaig, 1991; Li et al., 2004). The level of wheat grain dormancy partly depends on ABA sensitivity before and after the grain reaches physiological maturity (Gubler et al., 2005; Shu et al., 2016; Sun et al., 2005). One well-characterized positive regulator of ABA signalling, *ABI4*, was initially identified in screens for mutants exhibiting ABA-resistant germination (Finkelstein, 1994). *ABI4* is a member of the AP2/ERF transcription factor family and can activate or repress gene expression by binding to specific *cis*-elements in gene promoters via its APETALA 2 (AP2) DNA-binding domain (Wind et al., 2013). It has been documented that *ABI4* interacts with target genes to regulate seed dormancy and germination. For example, *ABI4*-dependent temporal regulation of *PTR2* expression influences water status during seed germination, promoting germination of imbibed grain (Choi et al., 2020). *ABI4* is indispensable for repressing the expression of *ARR6/7/15*, which is involved in seed dormancy (Huang et al., 2017). Moreover, *ABI4* is a primary positive regulator of *ABI4, ABI5*, and *SBE2.2*, activating transcription by binding the CACCG-box (CE1-like) in the promoter regions during seed development (Bossi et al., 2009). Apart from *ABI4* itself, various transcription factors could regulate *ABI4* transcription, including several *WRKY* transcription factors that can bind to the W-box sequence in the *ABI4* promoter region (Antoni et al., 2011; Liu et al., 2012; Shang et al., 2010). *Myb96* induces *ABI4* expression by binding to its promoter during seed germination and seedling development (Lee and Seo, 2015).

In addition to *ABI4*, two other transcription factors (*ABI3/VP1* and *ABI5*) have been characterized that regulate ABA response during seed development (Finkelstein, 1994; Finkelstein and Lynch, 2000). It has been reported that some cross-regulation of expression existed among *ABI3, ABI4*, and *ABI5*, whose function in a combinatorial network, rather than a regulatory hierarchy, controlling seed development and ABA response (Soderman et al., 2000). Moreover, *ABI3, ABI4, ABI5* have similar effects on seed dormancy and the expression of maturation-specific seed proteins (Finkelstein, 1994). However, *ABI4* is a focal point in the signal transduction pathways of ABA (Niu et al., 2002). Orthologues of *ABI4* have been reported in many other plant species, including maize, rice, and lotus (Ming et al., 2013; Niu et al., 2002; Wang et al., 2015). In maize, *ZmABI4* is seed-specific, reaching maximum expression at 20 days post-anthesis (DPA) (Niu et al., 2002). In the rice database, a single sequence shares significant homology with the *AtABI4* AP2 domain, indicating that a single *ABI4* homolog exists in rice (Yu et al., 2002). However, there is limited information available for *ABI4* orthologues in wheat.

Synthetic hexaploid wheat SHW-L1 obtained from the hybridization of *Triticum turgidum* and *Aegilops tauschii* is a useful genetic resource and shows significant tolerance to PHS (Yang et al., 2014). To investigate the regulatory factors that interact with *TaABI4* and the role of *TaABI4* in the ABA-induced seed dormancy pathway, we performed a conservation analysis on *ABI4* in wheat ancestral species and modern cultivars and subsequently cloned this gene. We analysed the expression pattern of *TaABI4* at different grain developmental stages. Furthermore, we carried out expression QTL analysis (eQTL) to detect regions regulating the expression of *TaABI4* in recombinant inbred lines (RILs), providing further insight into the role of *TaABI4* in ABA signal transduction pathways and into the regulatory framework that controls seed germination in wheat.

## Materials and methods

### Plant material

Chuanmai32 (CM32, PHS susceptible), synthetic hexaploid wheat (SHW-L1, PHS resistant), and their derived RILs (138 lines) were grown in glasshouse conditions (16-h light, 8-h dark, 22 °C, 70% relative humidity). Days to flowering was measured for each spikelet based on the anther extrusion at 50% of the spike. Developing grains from 5 days post-anthesis (DPA) to 30DPA were collected at five-day intervals from the centre florets for subsequent gene expression profiling. Young leaves of SHW-L1 and CM32 were used for DNA extraction. Each sample had biological replicates and was immediately frozen into liquid nitrogen and stored at −80°C for RNA extraction.

### Sequence characterization and in-silico promoter analysis

Based on the results of BLASTP searches, we obtained coding sequences of *TaABI4* in Chinese Spring using EnsemblPlants (http://plants.ensembl.org/index.html). Protein domains of genes were predicted using the SMART tool (http://smart.embl-heidelberg.de/). The coding sequences of *TaABI4* were used to query the target database (ViroBLAST, http://202.194.139.32/blast/viroblast.php, and The Wheat ‘Pan Genome’, http://www.10wheatgenomes.com/data-repository/, The Aegilops tauschii genome, http://aegilops.wheat.ucdavis.edu/ATGSP/data.php) to download homologous genes and 2 kb upstream sequences from translational initiation codon in 17 wheat cultivars and three wheat ancestors (Altschul et al., 1997; Ling et al., 2018; Luo et al., 2017; Zhu et al., 2019)(**Table S1**).

Amino acid sequences were aligned using DNAMAN (Version. 5.2.10, Lynnon Biosoft, Quebec, Canada). Putative *cis*-acting regulatory elements located in promoter regions were predicted using PLANTCARE (Lescot et al., 2002) (http://bioinformatics.psb.ugent.be/webtools/plantcare/html/) and PLACE (Higo et al., 1999) (http://www.dna.affrc.go.jp/PLACE/). The analysis of conserved motifs in 17 wheat cultivars was obtained using the MEME suite (Bailey et al., 2009) (http://meme-suite.org/tools/meme). This program was used to search for the top 5 *cis*-motifs with consensus patterns of 6~50 base width and E-value < 0.01, on the forward strand of the input sequences only.

### Prediction of proteins and PEST motifs

Generated coding sequences were translated to predicted proteins using DNAMAN with default parameters. Searches for potential PEST sequences were performed using the ePESTfind (http://www.bioinformatics.nl/cgi-bin/emboss/epestfind). We used the input parameters in all cases and defined that a score above zero denoted a possible PEST sequence (Gregorio et al., 2014).

### PCR amplification

According to *TaABI4* nucleotide sequences of Chinese Spring, specific primers for the gene were designed online (https://www.ncbi.nlm.nih.gov/tools/primer-blast/) and shown in **Table S2**. Genomic DNA was isolated from SHW-L1/CM32 young leaves using the CTAB method (Zhang et al., 2013) and was used as templates to amplify the DNA sequences of *TaABI4*. PCR was performed using high-fidelity Prime STAR Polymerase (TaKaRa, Japan) under the following conditions: 98 °C for 3 min; 35 cycles of 98 °C for 50 s, 60-65 °C for 50 s, and 72 °C for 90 s; followed by a final extension step of 72 °C for 10 min. The PCR amplification products were ligated into the pEASY-blunt Cloning Vector (TransGen, China), and the resulting ligation mixtures were transformed into *E.coli* Trans1-T1 chemically competent cells (TransGen, China) to obtain positive clones for sequencing.

### RNA extraction and expression analysis

Primer pairs in the relevant conserved exon regions of *TaABI4* among A, B, and D genomes in SHW-L1 and CM32 were used to amplify 151 bp amplicons (**Table S2**). The expression level of *TaABI4* was measured in the parents at six seed development stages (5, 10, 15, 20, 25, and 30DPA). RNA was extracted from each sample using the total RNA extraction kit (Biofit, China), and genomic DNA was removed with DNaseI.

Three seed-developing stages (10DAP, 20AP, 30DAP) of SHW-L1/CM32 were selected to carry out RNA sequencing (RNAseq). RNA quantity and quality were assessed using a NanoDrop 2000 spectrophotometer (Thermo Scientific, USA) and checked for integrity on an Agilent 2100 bioanalyser (Agilent Technologies, USA) by denaturing agarose gel electrophoresis with ethidium bromide staining. Equimolar amounts of the libraries were constructed and sequenced by BerryGenomics (Beijing) using the Illumina HiSeq-2000 and HiSeq X Ten platform (Illumina, USA). Gene transcript levels were estimated using transcripts per million (TPM) (Zhao et al., 2020).

First-strand cDNA was synthesized using a PrimeScriptTM 1st Strand cDNA Synthesis Kit (Takara). cDNA sampling was performed in duplicate and SsoFast™ EvaGreen^®^ Supermix (Bio-Rad) used for real-time quantitative PCR (RT-qPCR) (CFX96 Touch™ Real-Time PCR Detection System, Bio-Rad, USA). Each reaction contained approximately 50ng first-strand cDNA, 0.5 μL 10 μmol/L gene-specific primers, and 10 μL real-time PCR SYBR Green (TIANGEN, Beijing, China). Amplification conditions were: 5 min at 95°C, followed by 40 cycles of 30 s at 95°C, 30 s at 60°C, 40 s at 72°C, and a final extension of 10 min at 72°C. Seven 1/10 dilutions of the recombinant plasmid cDNA template were used to make a standard curve for amplification efficiency (E) calculation. Three housekeeping genes, *TaActin, Ta.14126.1*, and *Ta.7894.3.al_at*, were used as internal controls (Long et al., 2010). Gene expression data were analysed using the Bio-Rad CFX Manager (Bio-Rad) software. The expression profile of the target gene was normalized to that of the internal control genes, and the geometric mean was calculated. The relative gene expression quantity of each sample was calculated using the E^−ΔΔCt^ method (Pfaffl, 2001).

### *Expression QTL* (eQTL) *analysis*

In order to characterize regions that regulate *TaABI4* expression levels, we conducted eQTL mapping analysis within the RIL population using the previously constructed high-density genetic map (Yang et al., 2019). eQTL analysis was achieved using the WinQTLcart2.5 software (North Carolina State University, Raleigh, NC, USA) with the composite interval mapping (CIM) method (Wang and Basten, 2007). The analysis was implemented by setting the control parameters to model 6 (standard model), forward regression, 10-cm windows, and five makers as the control. The threshold was set at 4.0 to detect eQTLs. The wheat reference genome “Chinese Spring,” IWGSC RefSeq v1.0 (International Wheat Genome Sequencing Consortium 2018) was used to query marker positions using the blastn2.2.26+ package (Camacho et al. 2009).

## Results

### Sequence characterization of TaABI4

The DNA sequence of *ABI4* (AT2G40220) from *Arabidopsis* was used as a query sequence to carry out BLAST searches in EnsemblPlants. Three homologues of *TaABI4* were identified on the A, B, and D sub-genomes of 18 wheat cultivars (*TaABI4-1A, TaABI4-1B, TaABI4-1D*). All the 50 *TaABI4* sequences were found to be represented by a single exon. The coding sequences (CDS) of three homologues of Chinese Spring were conserved with 97.53% nucleotide identity. Compared to *TaABI4-1A, TaABI4-1B* and *TaABI4-1D* had two 3-6 bp deletions as well as 28 single-nucleotide polymorphisms (SNPs), 13 SNPs of which caused non-synonymous mutations (**Fig. S1**). The three homologues encoded proteins with 260, 256, and 257 amino acid residues, respectively. The proteins have highly conserved AP2 domains that were also found in previously annotated *AtABI4* in *Arabidopsis* and *ZmABI4* in maize (*Zea mays*) (**Fig. 1**). In addition, ten amino acids (KGGPENAKFR) were contiguous to the AP2 domain (designated as the AP2-associated motif). Additionally, a stretch of eight amino acids (LRPLLPRP) identified as the LRP motif was located nearby. (**Fig. 1**). TaABI4 proteins revealed 100% identity in these common regions, while TaABI4-1A contained three additional amino acids His_171_, Leu_196_, and Ala_197_ (**Fig. 1**). Putative proteins were predicted from *Ae. tauschii, T. dicoccoides* cv. Zavitan and *T. urartu*. The protein sequence of AetABI4 obtained from *Ae. tauschii* illustrated 100% identity with TaABI4-1D sequence. TuABI4-1A obtained from *T. urartu*, showed 99.23% amino acid identity with TaABI4-1A. TdABI4-1B of *T. dicoccoides* cv. Zavitan shared 98.08% identity with TaABI4-1B (**Fig. 1**).

**Figure 1.**
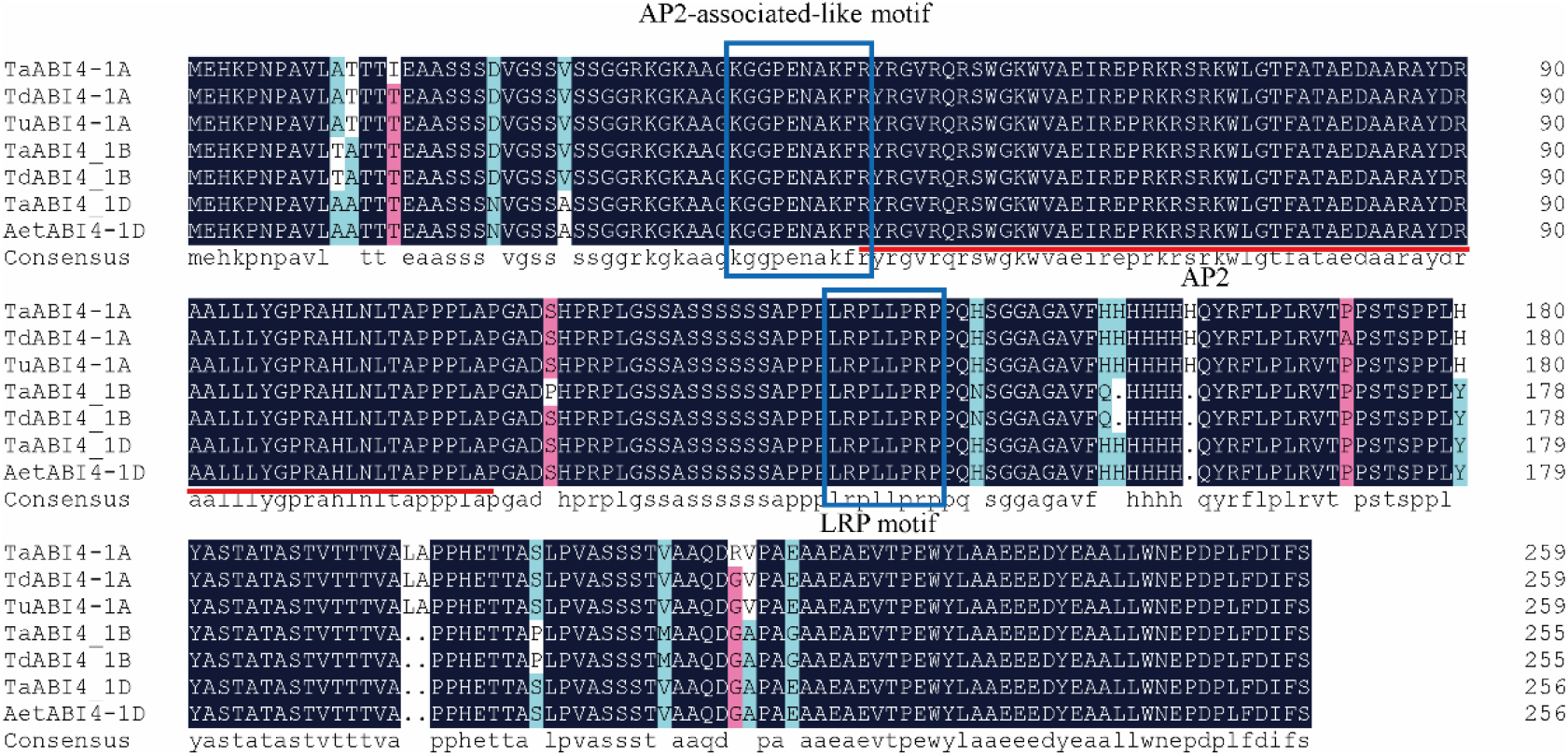
The alignment of *ABI4* proteins of Chinese Spring and wheat ancestors. Sequences including *TaABI4-1A/TaABI4-1B/TaABI4-1D* from Chinese Spring, *TdABI4-1A, and TdABI4-1B* from *T. dicoccoides* cv. Zavitan, *TuABI4-1A* from *T. Urartu*, and *AetABI4-1D from Ae. tauschii*. The AP2 region was indicated by red lines at the bottom, LRP and AP2-associated-like motifs were boxed.

### ABI4 proteins and putative motifs analysis in wheat cultivars

The AP2 domains, the AP2-associated motifs, and the LRP motifs were conserved in 50 putative ABI4 homologous proteins in terms of their position and sequence identity (**Fig. 2**). Putative PEST degradation signals at the terminus of wheat ABI4 proteins with a positive probability value (>0) were detected using the PEST-find program (Rice et al., 2000), which was in agreement with a previous report (Gregorio et al., 2014). It demonstrated that potential PEST sequences were detected in all of these proteins, with probability scores ranging from +0.44 (ABI4-1A) to +3.52 (ABI4-1B) (**Table 1**). For ABI4-1B and ABI4-1D proteins, one PEST sequence was detected at the C-terminal with a length of 60AA. For ABI4-1A proteins, a shorter PEST motif of 41 amino acids was detected, sharing 99.8% identity amongst the 17 cultivars in addition to another PEST sequence predicted at the N-terminus that was also identified in TaABI4-1B. Although some variant amino acids were detected in proteins of each genome, as shown in grey boxes in **Fig. 2**, they did not locate in the region of crucial motifs. It demonstrates that the ABI4 proteins are conserved in their protein architecture, coinciding with their central role in wheat hormone signalling.

**Table 1.**
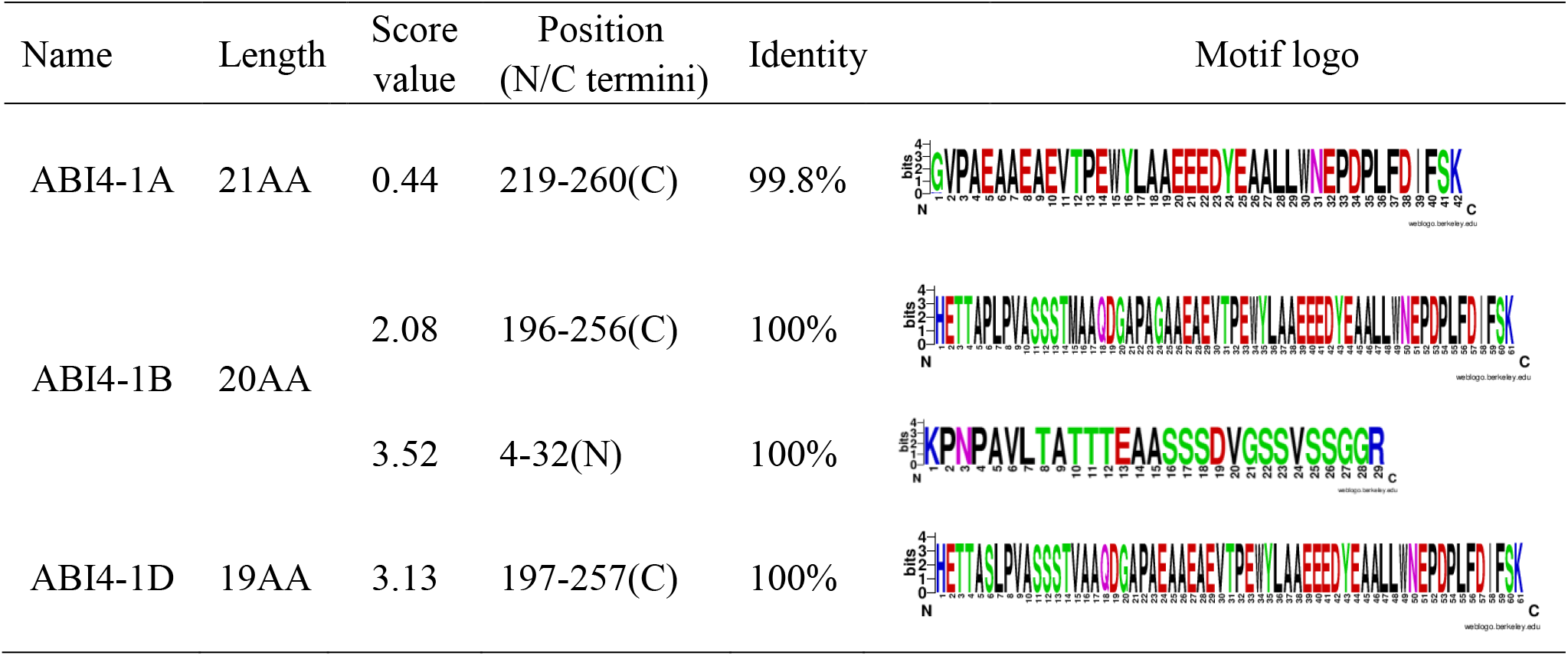
Conservation of putative PEST sequences in ABI4 proteins from wheat cultivars.

| Name | Length | Score value | Position (N/C termini) | Identity | Motif logo |
| --- | --- | --- | --- | --- | --- |
| ABI4-1A | 21AA | 0.44 | 219-260(C) | 99.8% |  |
| ABI4-1B | 20AA | 2.08 | 196-256(C) | 100% |  |
|  |  | 3.52 | 4-32(N) | 100% |  |
| ABI4-1D | 19AA | 3.13 | 197-257(C) | 100% |  |

**Figure 2.**
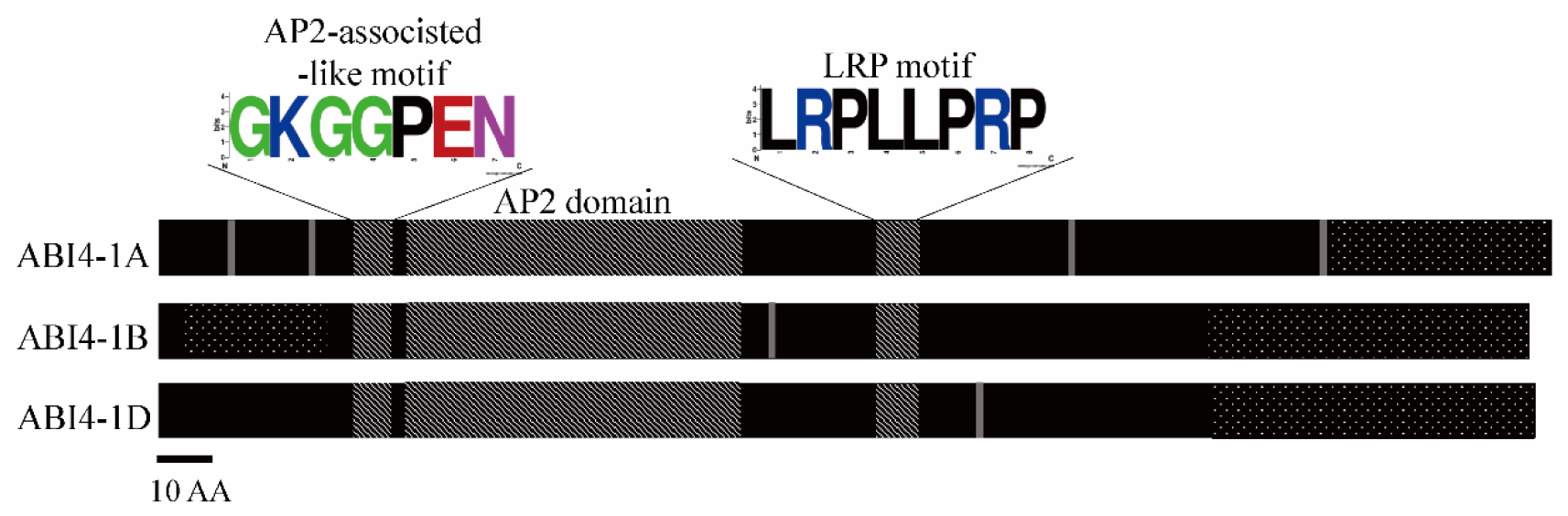
Protein structure schematic diagram of ABI4 in wheat cultivars. Grey boxes indicate polymorphism of amino acid sequences and black boxes were highly conserved Amino Acid sequences. Boxes filled with twill were conserved domains and motifs, and boxes filled with dots were potential PEST motifs.

### Potential cis-acting regulatory elements of ABI4 promoters in wheat ancestors

The presence of potential *cis*-regulatory elements at upstream (≥ 2,000 bp) region of *TaABI4* homologues from wheat cultivar Chinese Spring was analysed. Eleven types of potential *cis*-acting regulatory elements were identified in the upstream region (**Fig. 3**). This region was also isolated from *T. dicoccoides* cv. Zavitan, *T. urartu*, and *Ae. tauschii*. A putative TATA-box was detected 190 bp upstream of the start codon. A binding site (CE1-like motif, CACCGCCCC) was present immediately downstream from a putative W-box (TTGACY). In addition, RY-elements with CATGCATG involved in seed-specific regulation were predicted. ABRE elementsknown to be involved in ABA response, with CACGTG core motif, were recognized nearby the 5’ termini. ARE elements with an AAACCA core motif that are essential for the anaerobic induction also existed in all ABI4 proteins. Additionally, conserved motifs such as CAAT box, CAT box, and A box were detected. One Myb and one Myc element, known to be involved in ABA signalling (Lin, 2009), were predicted in *TaABI4-1D* and *AetABI4* promoter regions. The detected *cis*-acting regulatory elements were conserved among the wheat and its ancestral species.

**Figure 3.**
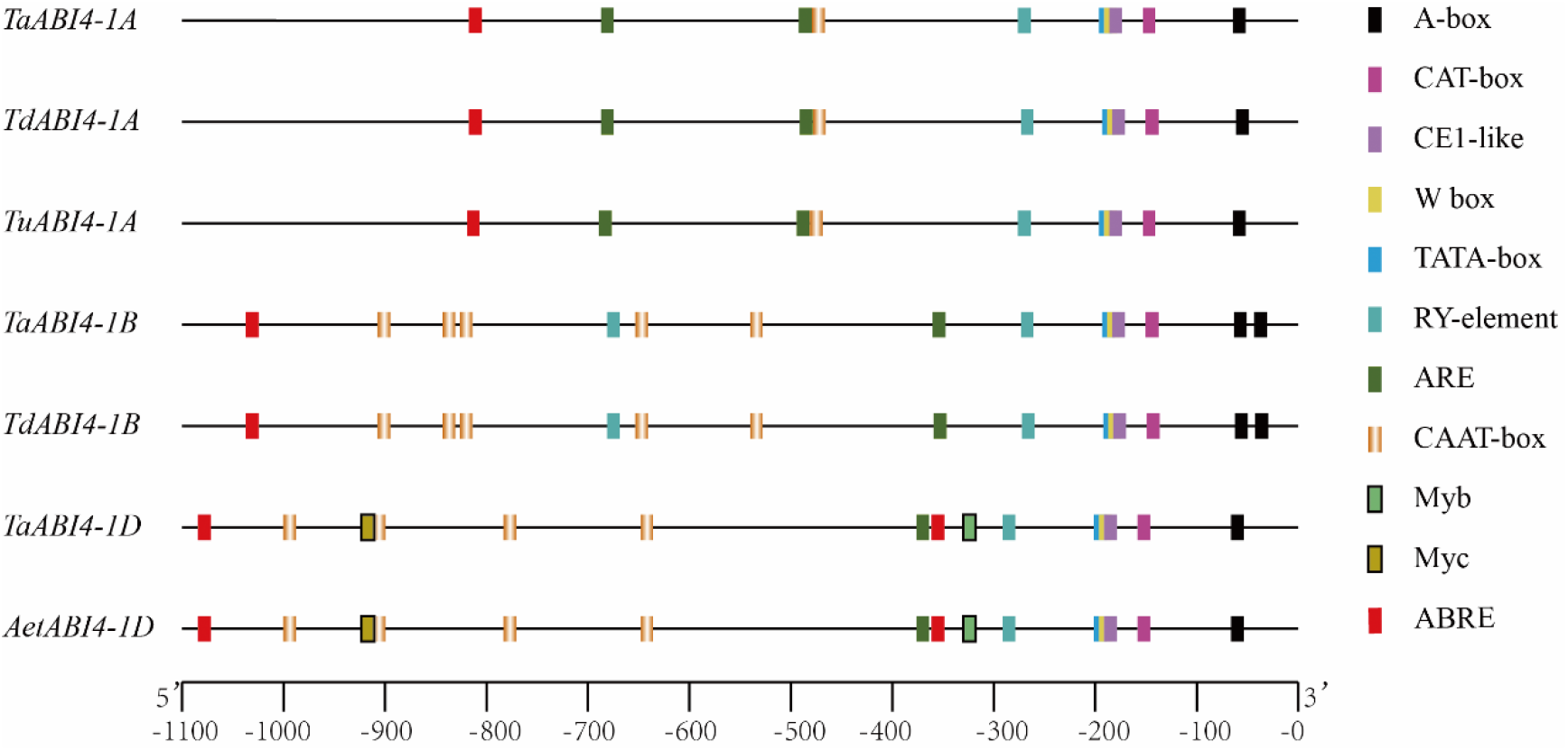
Potential *cis*-acting regulatory elements in upstream regions of *ABI4* genes from wheat and wheat ancestors. Colored boxes represent different *cis*-regulatory elements.

### The putative motifs analysis of ABI4 genes in wheat cultivars

The top five motifs identified by this analysis were found in almost all of the *ABI4* genes in wheat cultivars and were highly conserved in terms of number and position **(Fig. 4)**. Although motif two did not exist in the A sub-genome of Kronos, it shared 99.4% identity among 50 upstream regions and was regarded as a novel *cis*-motif with no current description in the PLACE database (**Table 2**). As shown in **Table 2**, motif 1 with a W-box as its core element was also conserved in all sequences with 100% identity. Although there were some variable SNPs in motif 3, motif 4, and motif 5, they did not exist in the core region of each motif. Overall, putative motifs within the upstream of *ABI4* genes were almost completely conserved in wheat cultivars.

**Table 2.**
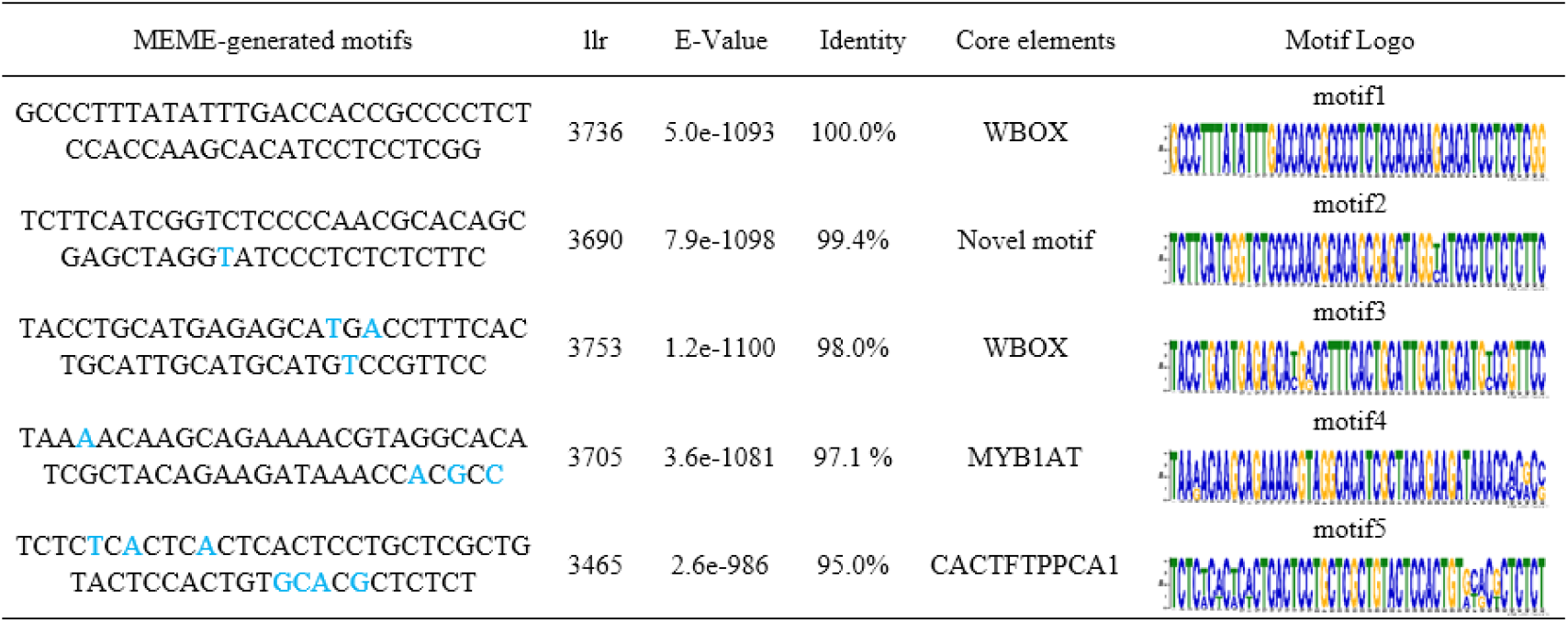
Conserved *cis*-motifs found in upstream promoter regions of *ABI4* genes in wheat cultivars. llr means log-likelihood ratio.

**Figure 4.**
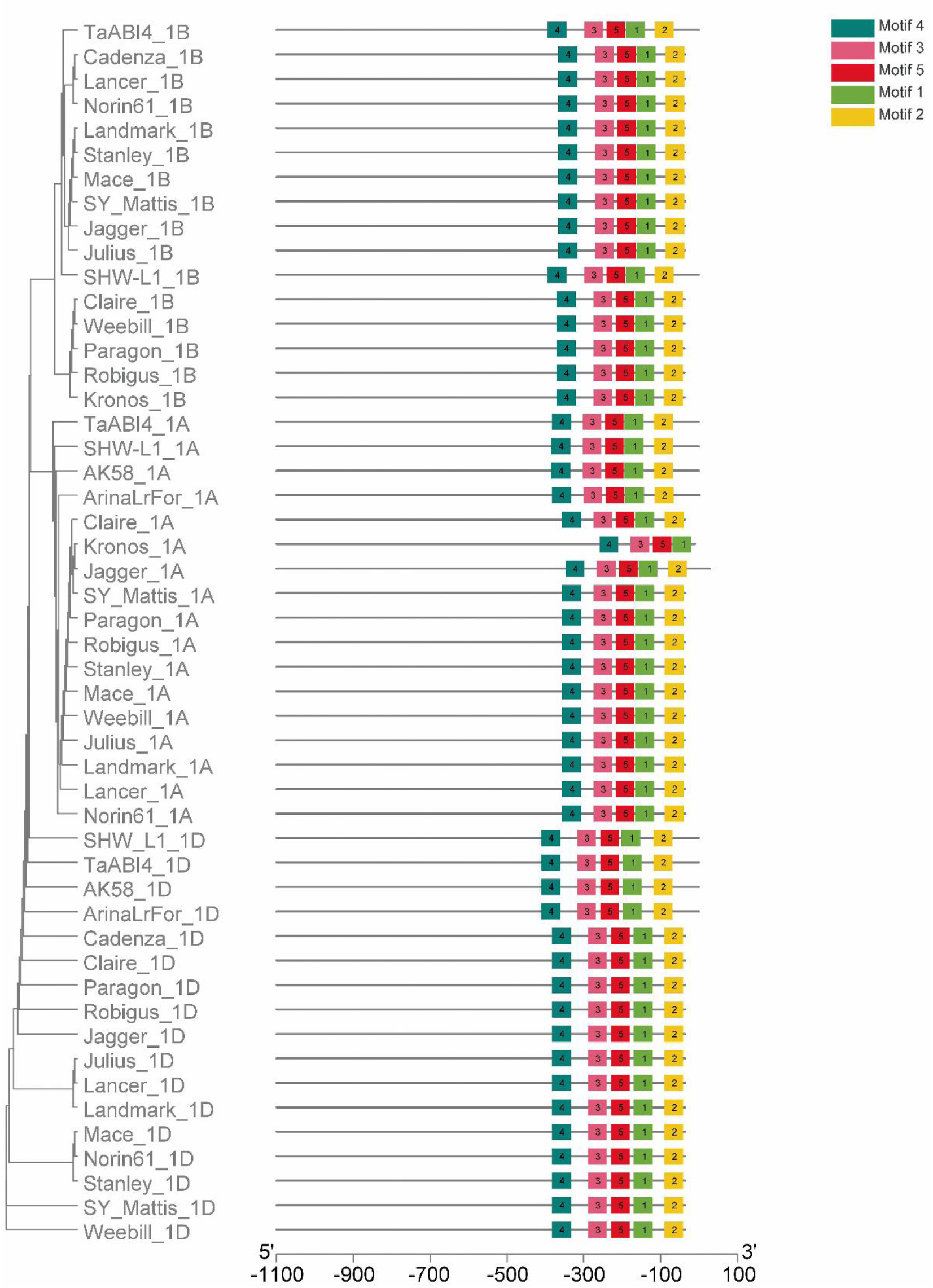
Schematic representation of conserved *cis*-motifs (obtained using MEME) in upstream regions of *ABI4* genes from wheat cultivars. Different motifs were represented by boxes of different colors.

### Cloning and qRT-PCR analysis of TaABI4 in SHW-L/CM32 developing seeds

The *TaABI4* sequences were cloned from SHW-L and CM32, which were highly conserved in these two cultivars (**Fig. S2**). According to RNAseq analysis, the expressional level of *TaABI4* in CM32 was higher than that in SHW-L1 at each detected stage (**Fig. 5A**). Then, RT-qPCR assays were performed using cDNA from five-time points (5, 15, 20, 25, and 30DPA) to detect expression level variation of *TaABI4* between SHW-L1 and CM32. During seed development, *TaABI4* expression began as early as 10DPA, increasing between 10DPA and 15DPA as the transition from growth to storage phase of grain development (starting after 12DPA) took place, and peaked at 20DPA with a decline in expression until 30DPA. The expression of *TaABI4* in CM32 was higher than that in SHW-L1 in most of the measured stages. The two most significant differences in relative expression were detected at 15DPA (5.07-fold) and 20DPA (1.39-fold) (**Fig. 5B**).

**Figure 5.**
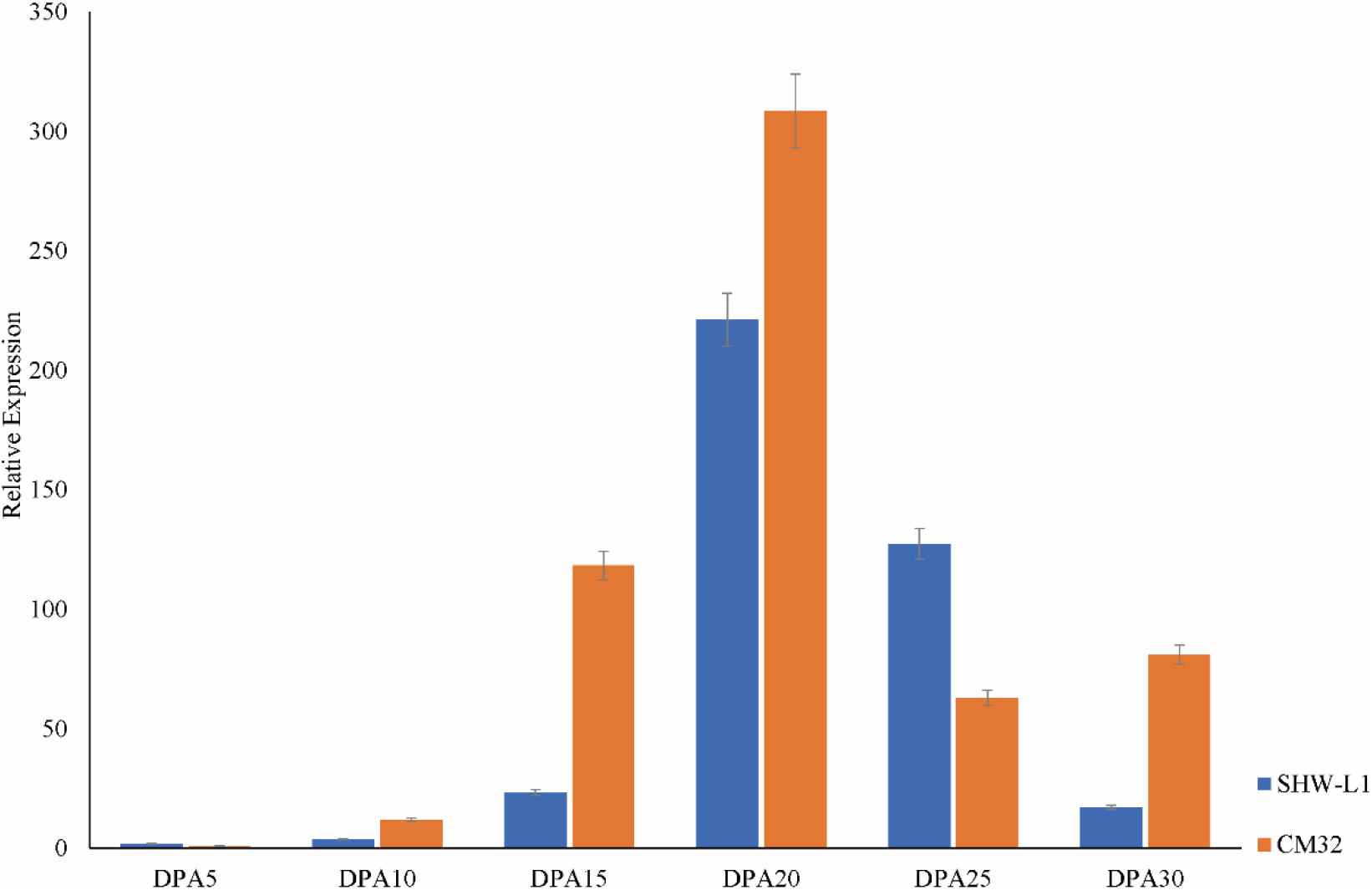
The expression pattern of *TaABI4* in SHW-L1 and CM32. (A) the expression assays using RNAseq. The y-axis denotes TPM (Transcripts Per Kilobase Million). (B) the expression assays using qRT-PCR.

### Expression QTL mapping

The significant difference between CM32 and SHW-L1 in expression levels of TaABI4 at 15 and 20DPA enabled the detection of eQTLs. Based on the consensus genetic map and corresponding SNP marker positions, six significant eQTLs (P < 0.05, LOD >4) were identified (Table 3; Fig.6). One eQTL detected on chromosome 2A at 15DAP was designated as *eQABI4.15DPA.2A.1*, with LOD scores at 4.53. Two eQTL regions located on chromosome 2D designated as *eQABI4.20DPA.2D.1* and *eQABI4.20DPA.2D.2* were detected at 20DAP, showing 9.63 and 6.38 LOD scores, respectively. *eQABI4.20DPA.4A.1* and *eQABI4.20DPA.4A.2* were located on chromosome 4D with negative alleles from SHW-L1, they explained 38.2% and 46.1% of the phenotypic variation, respectively. Physical mapping of 3B eQTL, designated as *eQABI4.20DPA.3B.1*, showing that the corresponding interval location was Chr.3B: 667902308-669428443. All identified eQTLs had negative additive effects, indicating that eQTLs that could decrease expression of *TaABI4* were derived from synthetic wheat SHW-L1.

**Table 3.**
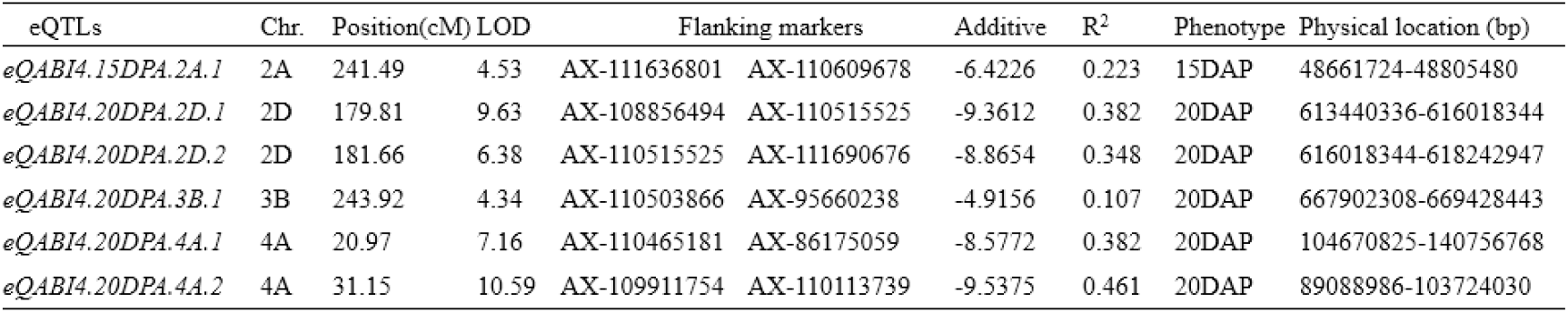
eQTL mapping results of *TaABI4* in SHW-L1 and CM32.

**Table 4.** Candidate genes expressed in seeds and ABA related genes near eQTL interval

| Gene ID | Orthologues |  | Description in Wheat Gmap |
| --- | --- | --- | --- |
|  | Species | Gene name |  |
| TraesCS2B02G600800 | *Ae. tauschii* | *GH3.3* | GH3 auxin-responsive promoter |
| TraesCS2B02G601300 | *Ae. tauschii* | *PUB23-like* | protein ubiquitination |
| TraesCS2B02G602000 | *Ae. tauschii* | *TMK1* | protein phosphorylation |
| TraesCS2B02G603000 | *A. thaliana* | *NHL12* | Late embryogenesis abundant protein, *LEA-14* |
| TraesCS2B02G603100 | *A. thaliana* | *NHL10* | Late embryogenesis abundant protein, *LEA-14* |
| TraesCS2B02G603200 | *A. thaliana* | *NHL10* | Late embryogenesis abundant protein, *LEA-14* |

**Figure 6.**
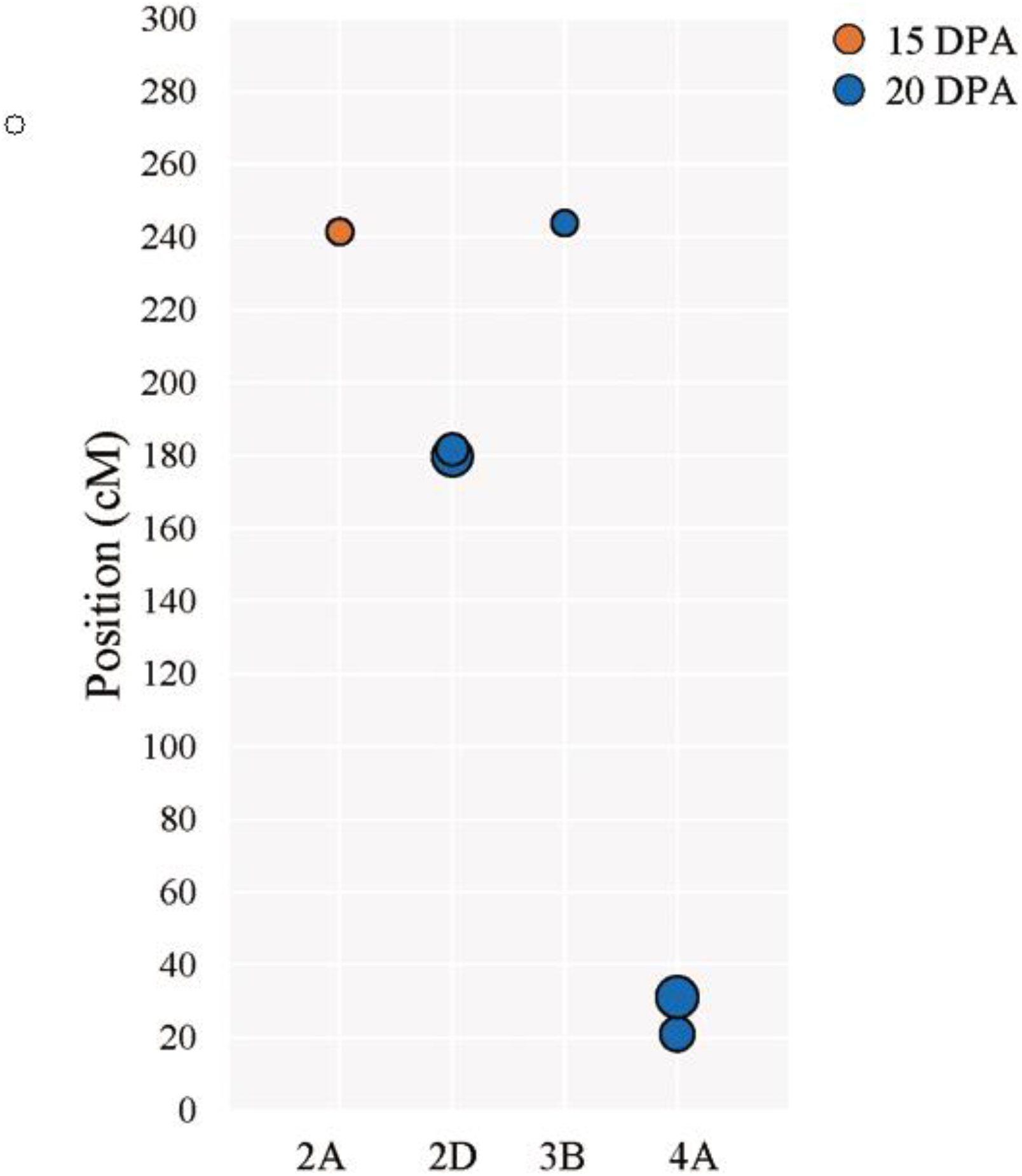
eQTL genetic locations in the genetic map. The size of the circles means LOD values. The x-axis denotes different chromosomes.

## Discussion

In this study, we presented the characterization of the wheat *ABI4*, a gene involved in ABA-responsiveness during seed development and germination. TaABI4 proteins from three wheat sub-genomes were conserved with an AP2 domain required for nuclear localization (AP2-associated motif), as well as regions for transcriptional activation (LRP motif). The conserved domains are used as hallmarks to identify ABI4 orthologues in different species (Gregorio et al., 2014). Although the protein sequences for the three homologues had slight polymorphisms, the overall identity was high (96.9%). Our results suggested that the AP2 proteins presented in wheat are the orthologues of the *Arabidopsis* ABI4 and should be considered as TaABI4-1A, TaABI4-1B, TaABI4-1D.

Compared with the ABI4 obtained from *T. urartu, Ae. tauschii* and *T. dicoccoides*, the amino acid variation existed only in TaABI4-1A (Thr_15_/Leu_15_, Gly_218_/Arg_218_) and TaABI4-1B (Ser_108_/Pro_108_) and were not located in core regulatory regions (**Fig. 1**). It was indicating that TaABI4 was highly conserved during the polyploidization and domestication processes of wheat. In *Arabidopsis*, the low accumulation of ABI4 resulted from both post-transcriptional and post-translational regulation (Finkelstein et al., 2011). PEST sequences are degradation motifs that can affect protein stability (Gregorio et al., 2014) and are characterized by regions enriched in the amino acid proline, glutamic acid, serine, and threonine (Rogers et al., 1986). Based on the available pan-genome data, we analysed 50 putative ABI4 proteins from 18 wheat cultivars to predict potential PEST motifs. Most of the possible PEST sequences were located in the N-terminal region of the protein and were longer than AtABI4 (**Table 1**). These differences may cause divergence in post-translational mechanisms compared with *Arabidopsis*. In fact, ZmABI4 also has two PEST motifs located in the N-terminus and C-terminus showing score values of +3.04 and +0.68, respectively (Gregorio et al., 2014).

The discovery of *cis*-acting regulatory elements in the promoter regions is essential to understanding the spatial and temporal expression patterns of *ABI4* genes. The six *cis*-acting regulatory elements were conserved in terms of position and sequence identify (**Fig. 3**). TATA-box is regarded as the core promoter element, and transcription factors bind to TATA-proximal regions (W-box, CE-1 like) having been shown to regulate downstream gene transcription (Busk et al., 1997; Heins et al., 1992; Phukan et al., 2016). Additionally, A-box and RY-element are *cis*-acting regulatory elements, and CAT-box is related to meristem expression in *Arabidopsis* (Sakata et al., 2010). ABRE (ABA-responsive elements) motifs are known to participate in response to ABA (Sarkar and Lahiri, 2013). *TaABI4-1D* contained two classical ABRE elements that are necessary to constitute an active ABA-responsive complex because a single ABRE is not sufficient to confer ABA-responsiveness (Ganguly et al., 2011; Hobo et al., 1999; Zhang et al., 2005). Identification of conserved *cis*-acting regulatory elements in *ABI4* promoters of wheat revealed that other transcription factors might regulate those homologues.

The expression pattern of *TaABI4* was variable between a modern wheat cultivar and synthetic wheat accession. It is noteworthy that *TaABI4* showed higher transcript accumulation in weakly dormant material (CM32) than in dormant material (SHW-L1) during most periods of seed development (**Fig. 5**). By contrast, seeds of the *Arabidopsis abi4* mutant germinated significantly more quickly than wild type (Shu et al., 2013), indicating the presence of functional *ABI4* is important for resistance to PHS. The expression of *TaABI3* and *TaABI5* in SHW-L1 were both significantly higher than those in CM32 (Zhou et al., 2016). These results are consistent with the corresponding research results in *Arabidopsis* and maize finding *ABI3* and *ABI5* are positive regulators of seed dormancy (Finkelstein and Lynch, 2000; Hoecker et al., 1995; McCarty et al., 1991). Gene expression patterns of *TaABI3* and *TaABI5* were similar to *TaABI4* in the early and middle stages of seed development (5~15DPA), signifying that *TaABI4* associating the ABA biosynthetic pathway with *TaABI3* and *TaABI5* as found in *Arabidopsis* (Lopez-Molina et al., 2002). From these results, other regulatory factors interacting with *TaABI4* are required to complete our understanding of the gene networks involving seed germination.

eQTLs mapping is an efficient approach to identify genetic loci controlling complex crop traits (Chen et al., 2010; Motomura et al., 2013). In this study, six significant eQTLs associated with *TaABI4* expression variation were identified on chromosomes 2A, 2D, 3B, and 4A (**Table 3**), suggesting that the observed differences in *TaABI4* expression in the RILs population were regulated in-part by *trans*-acting factors (Doss et al., 2005). Several previous studies mapped the major QTLs for seed dormancy and PHS tolerance to chromosomes 4A (Mares et al. 2005; Torada et al. 2005; Chen et al. 2008). In this study, two major eQTLs located on chromosome 4A accounted for 38.2 and 46.1% of the phenotypic variance. This result further confirmed that the eQTLs on chromosome 4A control the expression of *TaABI4* to regulate seed germination in wheat. Some QTLs associated with the PHS were also detected on chromosome 2D. For instance, *QPhs.cnl-2D.1* is mapped for PHS resistance in white wheat (Munkvold et al, 2009). These results suggest that these eQTL regions detected in this study may provide candidate genes that play potential roles in regulating PHS through effects on *TaABI4* expression. *TaABI5* that reported as pre-harvest sprouting resistance gene (Zhou et al., 2016) was located near the *eQABI4.20DPA.3B.1* (<30Mb), signifying that *TaABI5* might regulate the expression of *TaABI4* in SHW-L1 and CM32. Thus, new eQTLs detected in this study suggested that unidentified genes or indirect regulation genes would affect *TaABI4*, which causes the different expression patterns of *TaABI4* compared with *Arabidopsis*.

In this study, the characterization of *TaABI4*, including its conserved protein domains and *cis*-acting regulatory elements analysis, provides information on the critical nucleotide and amino acid residues of this gene. Meanwhile, high conservation in amino acid sequences and promoter regions, but the different expression level of *TaABI4* in two wheat cultivars drove us to identify regions linked to candidate genes that function upstream of *TaABI4* transcripts. Six potential eQTL regions that may regulate the expression of *TaABI4* were detected. These results can be utilized for future *TaABI4* studies on interactions with other transcription factors in response to ABA and establishment of the co-expressed networks relating to seed germination, which will successfully boost the efficiency of wheat breeding with sufficient seed dormancy to prevent PHS.

## Acknowledgments

We thank Dr. Jizeng Jia from the Chinese Academy of Agricultural Science to provide the data of Aikang58.

## Declare of conflict interest

The authors declare no conflicts of interest.

## Supplementary Materials

***Table S1***. Currently available wheat genome assemblies for varieties different to the reference Chinese Spring landrace and wheat ancestors. (file type: MS Word document; file size: 19kb)

***Table S2*.** Primers used for amplification and expressional profile assay of *TaABI4*. (file type: MS Word document; file size: 16kb)

***Figure S1*.** The alignment of coding sequences including *TaABI4-1A, TaABI4-1B* and *TaABI4-1D* in Chinese Spring. (file type: PDF; file size: 58kb)

***Figure S2*.** The alignment of coding sequences including *TaABI4-1A, TaABI4-1B* and *TaABI4-1D* in SHW-L1 and CM32. (file type: PDF; file size: 77kb)

