## Supplementary material for "Characterization and expression QTL analysis of *TaABI4*, a pre-harvest sprouting related gene in wheat": TABLE S1

**Table S1.** Currently available wheat genome assemblies for varieties different to the reference Chinese Spring landrace and wheat ancestors.

| Wheat ancestor/cultivar | Database | Genome | Region |
| --- | --- | --- | --- |
| *T.urartu* | ViroBLAST | *Triticum urartu* cultivar G1812 | - |
| *T.dicoccoides cv.* Zavitan | ViroBLAST | 151210_zavitan_v2_pseudomolecules | - |
| *Ae. tauschii* | The *Aegilops tauschii* genome | Aet v4.0 | - |
| Jagger | The Wheat 'Pan Genome'  The Wheat 'Pan Genome'  The Wheat 'Pan Genome'  The Wheat 'Pan Genome'  The Wheat 'Pan Genome'  The Wheat 'Pan Genome'  The Wheat 'Pan Genome'  The Wheat 'Pan Genome'  The Wheat 'Pan Genome'  The Wheat 'Pan Genome'  ViroBLAST  ViroBLAST  ViroBLAST  ViroBLAST  ViroBLAST | 180529_Jagger_pseudomolecule_v1.1 | USA |
| Julius |  | 170807_Julius_MAGIC3_pseudomolecules | Germany |
| Lancer |  | 181120_Lancer_pseudomolecule_v1.0 | Australia |
| Landmark |  | 170831_Landmark_pseudomolecule | Canada |
| Mace |  | 181120_Mace_pseudomolecule_v1.0 | Australia |
| Norin61 |  | 190307_Norin61_pseudomolecule_v1.1 | Japan |
| Stanley |  | 180902_Stanley_pseudomolecules_v1.2 | Canada |
| SY_Mattis |  | 181016_SY_Mattis_pseudomolecule_v1.0 | France |
| Weebill |  | Weebill_1_NIAB_EI_scaffolds_v1.0 | Mexico |
| ArinaLrFor |  | 180808_Arina_pseudomolecules_v3 | Switzerland |
| Cadenza |  | *Triticum aestivum*_Cadenza_EIv1.1 | UK |
| Claire |  | *Triticum aestivum*_Claire_EIv1.1 | UK |
| Paragon |  | *Triticum aestivum*_Claire_EIv1.1 | UK |
| Robigus |  | *Triticum aestivum*_Robigus_EIv1.1 | UK |
| Kronos |  | *Triticum turgidum*_Kronos_EIv1.1 | UK |
| SHW-L1 | NCBI | PRJNA605937 | China |
| AK58* |  |  | China |

***** Dr. Jizeng Jia from the Chinese Academy of Agricultural Science to provide the data.
