## Supplementary material for "Characterization and expression QTL analysis of *TaABI4*, a pre-harvest sprouting related gene in wheat": FIGURE S2

```
SHW-L1_TaABI4-1A   ATGGAACACAAACCCAACCCAGCAGTGCTTGCCACTACCACTATCGAGGCCGCAAGCAGCAGCGACGTCGGGAGCAGCGTGAGCAGCGGCGGCCGGAAGG   100
CM32_TaABI4-1A     ATGGAACACAAACCCAACCCAGCAGTGCTTGCCACTACCACTATCGAGGCCGCAAGCAGCAGCGACGTCGGGAGCAGCGTGAGCAGCGCCGGCCGGAAGG   100
SHW-L_TaABI4-1B    ATGGAACACAAACCCAACCCAGCAGTGCTTACCGCTACCACTACCGAGGCCGCAAGCAGCAGCGACGTCGGGAGCAGCGTGAGCAGTGGCGGCCGGAAGG   100
CM32_TaABI4-1B     ATGGAACACAAACCCAACCCAGCAGTGCTTACCGCTACCACTACCGAGGCCGCAAGCAGCAGCGACGTCGGGAGCAGCGTGAGCAGTGGCGGCCGGAAGG   100
SHW-L1_TaABI4-1D   ATGGAACACAAACCCAACCCAGCAGTGCTTGCCGCTACCACTACCGAGGCCGCAAGCAGCAGCAACGTCGGGAGCAGCGCGAGCAGCGGCGGCCGGAAGG   100
CM32_TaABI4-1D     ATGGAACACAAACCCAACCCAGCAGTGCTTGCCGCTACCACTACCGAGGCCGCAAGCAGCAGCAACGTCGGGAGCAGCGCGAGCAGCGGCGGCCGGAAGG   100
Consensus          atggaacacaaacccaacccagcagtgctt cc ctaccacta cgaggccgcaagcagcagc acgtcgggagcagcg gagcag g cggccggaagg

SHW-L1_TaABI4-1A   GGAAGGCGGCGGGCAAGGGCGGGCCGGAGAACGCCAAGTTCAGGTACCGTGGCGTGCGGCAGCGGAGCTGGGGCAAGTGGGTGGCGGAGATCCGGGAGCC   200
CM32_TaABI4-1A     GGAAGGCGGCGGGCAAGGGCGGGCCGGAGAACGCCAAGTTCAGGTACCGTGGCGTGCGGCAGCGGAGCTGGGGCAAGTGGGTGGCGGAGATCCGGGAGCC   200
SHW-L_TaABI4-1B    GGAAGGCGGCGGGCAAGGGCGGGCCGGAGAACGCCAAGTTCAGGTACCGTGGCGTGCGGCAGCGGAGCTGGGGCAAGTGGGTGGCGGAGATCCGGGAGCC   200
CM32_TaABI4-1B     GGAAGGCGGCGGGCAAGGGCGGGCCGGAGAACGCCAAGTTCAGGTACCGTGGCGTGCGGCAGCGGAGCTGGGGCAAGTGGGTGGCGGAGATCCGGGAGCC   200
SHW-L1_TaABI4-1D   GGAAGGCGGCGGGCAAGGGCGGGCCGGAGAACGCCAAGTTCAGGTACCGTGGCGTGCGGCAGCGGAGCTGGGGCAAGTGGGTGGCGGAGATCCGGGAGCC   200
CM32_TaABI4-1D     GGAAGGCGGCGGGCAAGGGCGGGCCGGAGAACGCCAAGTTCAGGTACCGTGGCGTGCGGCAGCGGAGCTGGGGCAAGTGGGTGGCGGAGATCCGGGAGCC   200
Consensus          ggaaggcggcgggcaagggcgggccggagaacgccaagttcaggtaccgtggcgtgcggcagcggagctggggcaagtgggtggcggagatccgggagcc

SHW-L1_TaABI4-1A   CCGCAAGCGCTCTCGCAAGTGGCTCGGCACCTTCGCCACTGCCGAGGACGCCGCGCGCGCCTACGACCGCGCGGCGCTGCTCCTCTACGGCCCGCGTGCC   300
CM32_TaABI4-1A     CCGCAAGCGCTCTCGCAAGTGGCTCGGCACCTTCGCCACTGCCGAGGACGCCGCGCGCGCCTACGACCGCGCGGCGCTGCTCCTCTACGGCCCGCGTGCC   300
SHW-L_TaABI4-1B    CCGCAAGCGCTCGCGCAAGTGGCTCGGCACCTTCGCCACTGCCGAGGATGCCGCGCGCGCCTACGACCGCGCGGCGCTGCTCCTCTACGGCCCACGTGCC   300
CM32_TaABI4-1B     CCGCAAGCGCTCGCGCAAGTGGCTCGGCACCTTCGCCACTGCCGAGGATGCCGCGCGCGCCTACGACCGCGCGGCGCTGCTCCTCTACGGCCCACGTGCC   300
SHW-L1_TaABI4-1D   CCGCAAGCGCTCGCGCAAGTGGCTCGGCACCTTCGCCACTGCCGAGGACGCCGCGCGCGCCTACGACCGCGCAGCGCTGCTCCTCTACGGCCCGCGTGCC   300
CM32_TaABI4-1D     CCGCAAGCGCTCGCGCAAGTGGCTCGGCACCTTCGCCACTGCCGAGGACGCCGCGCGCGCCTACGACCGCGCAGCGCTGCTCCTCTACGGCCCGCGTGCC   300
Consensus          ccgcaagcgctc cgcaagtggctcggcaccttcgccactgccgagga gccgcgcgcgcctacgaccgcgc gcgctgctcctctacggccc cgtgcc

SHW-L1_TaABI4-1A   CACCTCAACCTCACCGCGCCGCCGCCCCTGGCCCCCGGGGCGGACTCCCACCCCCGGCCCTTGGGGTCCTCGGCTTCTTCCTCGAGCTCCTCCGCGCCTC   400
CM32_TaABI4-1A     CACCTCAACCTCACCGCGCCGCCGCCCCTGGCCCCCGGGGCGGACTCCCACCCCCGGCCCTTGGGGTCCTCGGCTTCTTCCTCGAGCTCCTCCGCGCCTC   400
SHW-L_TaABI4-1B    CACCTCAACCTCACCGCGCCGCCGCCCCTGGCCCCCGGGGCGGACCCCCACCCCCGGCCCTTGGGGTCCTCGGCTTCTTCCTCGAGCTCCTCCGCGCCCC   400
CM32_TaABI4-1B     CACCTCAACCTCACCGCGCCGCCGCCCCTGGCCCCCGGGGCGGACCCCCACCCCCGGCCCTTGGGGTCCTCGGCTTCTTCCTCGAGCTCCTCCGCGCCCC   400
SHW-L1_TaABI4-1D   CACCTCAACCTCACCGCGCCGCCGCCCCTGGCCCCCGGGGCGGACTCCCACCCCCGGCCCTTGGGGTCCTCGGCTTCTTCCTCGAGCTCCTCCGCGCCCC   400
CM32_TaABI4-1D     CACCTCAACCTCACCGCGCCGCCGCCCCTGGCCCCCGGGGCGGACTCCCACCCCCGGCCCTTGGGGTCCTCGGCTTCTTCCTCGAGCTCCTCCGCGCCCC   400
Consensus          cacctcaacctcaccgcgccgccgcccctggcccccggggcggac cccacccccggcccttggggtcctcggcttcttcctcgagctcctccgcgcc c

SHW-L1_TaABI4-1A   CGCCGCTCCGGCCGCTCTTGCCGCGCCCGCCGCAGCACTCAGGTGGCGCCGGGGCGGTCTTCCACCACCACCACCACCACCACCAATACCGCTTCCTGCC   500
CM32_TaABI4-1A     CGCCGCTCCGGCCGCTCTTGCCGCGCCCGCCGCAGCACTCAGGTGGCGCCGGGGCGGTCTTCCACCACCACCACCACCACCACCAATACCGCTTCCTGCC   500
SHW-L_TaABI4-1B    CGCCGCTCCGGCCGCTCTTGCCGCGCCCGCCGCAGAACTCAGGTGGCGCCGGGGCGGTCTTCCAACACCACCACCAC......CAATACCGCTTCCTGCC   494
CM32_TaABI4-1B     CGCCGCTCCGGCCGCTCTTGCCGCGCCCGCCGCAGAACTCAGGTGGCGCCGGGGCGGTCTTCCAACACCACCACCAC......CAATACCGCTTCCTGCC   494
SHW-L1_TaABI4-1D   CGCCGCTCCGGCCGCTCTTGCCGCGCCCGCCGCAGCACTCAGGTGGCGCCGGGGCGGTCTTCCAACACCACCACCACCAC...CAATACCGCTTCCTGCC   497
CM32_TaABI4-1D     CGCCGCTCCGGCCGCTCTTGCCGCGCCCGCCGCAGCACTCAGGTGGCGCCGGGGCGGTCTTCCACCACCACCACCACCAC...CAATACCGCTTCCTGCC   497
Consensus          cgccgctccggccgctcttgccgcgcccgccgcag actcaggtggcgccggggcggtcttcca caccaccaccac      caataccgcttcctgcc

SHW-L1_TaABI4-1A   GCTCCGCGTGACTGCACCGTCCACGTCGCCGCCTCTGCACTACGCGAGCACAGCCACCGCGTCCACGGTGACCACCACGGTGGCGCTGGCGCCGCCGCAC   600
CM32_TaABI4-1A     GCTCCGCGTGACTGCACCGTCCACGTCGCCGCCTCTGCACTACGCGAGCACAGCCACCGCGTCCACGGTGACCACCACGGTGGCGCTGGCGCCGCCGCAC   600
SHW-L_TaABI4-1B    GCTCCGTGTGACGCCACCGTCGACGTCGCCGCCTCTGTACTACGCGAGCACAGCCACCGCGTCCACGGTGACCACCACGGTGGCGC......CGCCTCAC   588
CM32_TaABI4-1B     GCTCCGTGTGACGCCACCGTCGACGTCGCCGCCTCTGTACTACGCGAGCACAGCCACCGCGTCCACGGTGACCACCACGGTGGCGC......CGCCTCAC   588
SHW-L1_TaABI4-1D   GCTCCGCGTGACGCCACCGTCCACGTCGCCGCCTCTGTACTACGCGAGCACAGCCACCGCGTCCACGGTGACCACGACGGTGGCGC......CGCCGCAC   591
CM32_TaABI4-1D     GCTCCGCGTGACGCCACCGTCCACGTCGCCGCCTCTGTACTACGCGAGCACAGCCACCGCGTCCACGGTGACCACAACGGTGGCGC......CGCCGCAC   591
Consensus          gctccg gtgac  caccgtc acgtcgccgcctctg actacgcgagcacagccaccgcgtccacggtgaccac acggtggcgc      cgcc cac

SHW-L1_TaABI4-1A   GAGACGACTGCCTCGTTACCAGAGGCTTCGTCATCTACGGTGGCCGCCCAGGATGGTGTGCCGGCGGAGGCGGCAGAGGCTGAGGTGACCCCCGAATGGT   700
CM32_TaABI4-1A     GAGACGACTGCCTCGTTACCAGTGGCTTCGTCATCTACGGTGGCCGCCCAGGATGGTGTGCCGGCGGAGGCGGCAGAGGCTGAGGTGACCCCCGAATGGT   700
SHW-L_TaABI4-1B    GAGACGACTGCCCCGTTGCCAGTGGCTTCGTCATCTACGATGGCCGCCCAGGATGGTGCTCCGGCGGGGGCGGCAGAGGCTGAGGTGACCCCCGAATGGT   688
CM32_TaABI4-1B     GAGACGACTGCCCCGTTGCCAGTGGCTTCGTCATCTACGATGGCCGCCCAGGATGGTGCTCCGGCGGGGGCGGCAGAGGCTGAGGTGACCCCCGAATGGT   688
SHW-L1_TaABI4-1D   GAGACGACTGCCTCGTTGCCAGTGGCTTCGTCATCTACGGTGGCCGCCCAGGATGGTGCGCCGGCGGAGGCGGCAGAGGCTGAGGTGACCCCCGAATGGT   691
CM32_TaABI4-1D     GAGACGACTGCCTCGTTGCCAGTGGCTTCGTCATCTACGGTGGCCGCCCAGGATGGTGCGCCGGCGGAGGCGGCAGAGGCTGAGGTGACCCCCGAATGGT   691
Consensus          gagacgactgcc cgtt ccag ggcttcgtcatctacg tggccgcccaggatggtg  ccggcgg ggcggcagaggctgaggtgacccccgaatggt

SHW-L1_TaABI4-1A   ACCTTGCCGCCGAGGAGGAGGACTACGAGGCGGCGCTGCTGTGGAATGAACCTGATCCCTTGTTCGACATCTTCTCCAAGTG   782
CM32_TaABI4-1A     ACCTTGCCGCCGAGGAGGAGGACTACGAGGCGGCGCTGCTGTGGAATGAACCTGATCCCTTGTTCGACATCTTCTCCAAGTG   782
SHW-L_TaABI4-1B    ACCTTGCCGCCGAGGAGGAGGACTACGAGGCCGCGCTGCTGTGGAATGAACCTGATCCCTTGTTCGACATCTTCTCCAAGTG   770
CM32_TaABI4-1B     ACCTTGCCGCCGAGGAGGAGGACTACGAGGCCGCGCTGCTGTGGAATGAACCTGATCCCTTGTTCGACATCTTCTCCAAGTG   770
SHW-L1_TaABI4-1D   ACCTTGCCGCCGAGGAGGAGGACTACGAGGCGGCGCTGCTGTGGAATGAACCTGATCCCTTGTTCGACATCTTCTCCAAGTG   773
CM32_TaABI4-1D     ACCTTGCCGCCGAGGAGGAGGACTACGAGGCGGCGCTGCTGTGGAATGAACCTGATCCCTTGTTCGACATCTTCTCCAAGTG   773
Consensus          accttgccgccgaggaggaggactacgaggc gcgctgctgtggaatgaacctgatcccttgttcgacatcttctccaagtg
```
