## Supplementary material for "Characterization and expression QTL analysis of *TaABI4*, a pre-harvest sprouting related gene in wheat": FIGURE S1

```
TaABI4-1A   ATGGAACACAAACCCAACCCAGCAGTGCTTGCCACTACCACTATCGAGGCCGCAAGCAGCAGCGACGTCGGGAGCAGCGTGAGCAGCGGCGGCCGGAAGG   100
TaABI4-1B   ATGGAACACAAACCCAACCCAGCAGTGCTTACCGCTACCACTACCGAGGCCGCAAGCAGCAGCGACGTCGGGAGCAGCGTGAGCAGTGGCGGCCGGAAGG   100
TaABI4-1D   ATGGAACACAAACCCAACCCAGCAGTGCTTGCCGCTACCACTACCGAGGCCGCAAGCAGCAGCAACGTCGGGAGCAGCGCGAGCAGCGGCGGCCGGAAGG   100
Consensus   atggaacacaaacccaacccagcagtgctt cc ctaccacta cgaggccgcaagcagcagc acgtcgggagcagcg gagcag ggcggccggaagg

TaABI4-1A   GGAAGGCGGCGGGCAAGGGCGGGCCGGAGAACGCCAAGTTCAGGTACCGTGGCGTGCGGCAGCGGAGCTGGGGCAAGTGGGTGGCGGAGATCCGGGAGCC   200
TaABI4-1B   GGAAGGCGGCGGGCAAGGGCGGGCCGGAGAACGCCAAGTTCAGGTACCGTGGCGTGCGGCAGCGGAGCTGGGGCAAGTGGGTGGCGGAGATCCGGGAGCC   200
TaABI4-1D   GGAAGGCGGCGGGCAAGGGCGGGCCGGAGAACGCCAAGTTCAGGTACCGTGGCGTGCGGCAGCGGAGCTGGGGCAAGTGGGTGGCGGAGATCCGGGAGCC   200
Consensus   ggaaggcggcgggcaagggcgggccggagaacgccaagttcaggtaccgtggcgtgcggcagcggagctggggcaagtgggtggcggagatccgggagcc

TaABI4-1A   CCGCAAGCGCTCTCGCAAGTGGCTCGGCACCTTCGCCACTGCCGAGGACGCCGCGCGCGCCTACGACCGCGCGGCGCTGCTCCTCTACGGCCCGCGTGCC   300
TaABI4-1B   CCGCAAGCGCTCGCGCAAGTGGCTCGGCACCTTCGCCACTGCCGAGGATGCCGCGCGCGCCTACGACCGCGCGGCGCTGCTCCTCTACGGCCCACGTGCC   300
TaABI4-1D   CCGCAAGCGCTCGCGCAAGTGGCTCGGCACCTTCGCCACTGCCGAGGACGCCGCGCGCGCCTACGACCGCGCAGCGCTGCTCCTCTACGGCCCGCGTGCC   300
Consensus   ccgcaagcgctc cgcaagtggctcggcaccttcgccactgccgagga gccgcgcgcgcctacgaccgcgc gcgctgctcctctacggccc cgtgcc

TaABI4-1A   CACCTCAACCTCACCGCGCCGCCGCCCCTGGCCCCCGGGGCGGACTCCCACCCCCGGCCCTTGGGGTCCTCGGCTTCTTCCTCGAGCTCCTCCGCGCCTC   400
TaABI4-1B   CACCTCAACCTCACCGCGCCGCCGCCCCTGGCCCCCGGGGCGGACCCCCACCCCCGGCCCTTGGGGTCCTCGGCTTCTTCCTCGAGCTCCTCCGCGCCCC   400
TaABI4-1D   CACCTCAACCTCACCGCGCCGCCGCCCCTGGCCCCCGGGGCGGACTCCCACCCCCGGCCCTTGGGGTCCTCGGCTTCTTCCTCGAGCTCCTCCGCGCCCC   400
Consensus   cacctcaacctcaccgcgccgccgcccctggcccccggggcggac cccacccccggcccttggggtcctcggcttcttcctcgagctcctccgcgcc c

TaABI4-1A   CGCCGCTCCGGCCGCTCTTGCCGCGCCCGCCGCAGCACTCAGGTGGCGCCGGGGCGGTCTTCCACCACCACCACCACCACCAATACCGCTTCCTGCC   500
TaABI4-1B   CGCCGCTCCGGCCGCTCTTGCCGCGCCCGCCGCAGAACTCAGGTGGCGCCGGGGCGGTCTTCCAACACCACCACCA......CCAATACCGCTTCCTGCC   494
TaABI4-1D   CGCCGCTCCGGCCGCTCTTGCCGCGCCCGCCGCAGCACTCAGGTGGCGCCGGGGCGGTCTTCCACCACCACCACCACCA...CCAATACCGCTTCCTGCC   497
Consensus   cgccgctccggccgctcttgccgcgcccgccgcag actcaggtggcgccggggcggtcttcca caccaccacca      ccaataccgcttcctgcc

TaABI4-1A   GCTCCGCGTGACTCCACCGTCCACGTCGCCGCCTCTGCACTACGCGAGCACAGCCACCGCGTCCACGGTGACCACCACGGTGGCGCTGGCGCCGCCGCAC   600
TaABI4-1B   GCTCCGTGTGACGCCACCGTCGACGTCGCCGCCTCTGTACTACGCGAGCACAGCCACCGCGTCCACGGTGACCACCACGGT......GGCGCCGCCTCAC   588
TaABI4-1D   GCTCCGCGTGACGCCACCGTCCACGTCGCCGCCTCTGTACTACGCGAGCACAGCCACCGCGTCCACGGTGACCACAACGGT......GGCGCCGCCGCAC   591
Consensus   gctccg gtgac ccaccgtc acgtcgccgcctctg actacgcgagcacagccaccgcgtccacggtgaccac acggt      ggcgccgcc cac

TaABI4-1A   GAGACGACTGCCTCGTTACCAGTGGCTTCGTCATCTACGGTGGCCGCCCAGGATCGTGTGCCGGCGGAGGCGGCAGAGGCTGAGGTGACCCCCGAATGGT   700
TaABI4-1B   GAGACGACTGCCCCGTTGCCAGTGGCTTCGTCATCTACGATGGCCGCCCAGGATGGTGCTCCGGCGGGGGCGGCAGAGGCTGAGGTGACCCCCGAATGGT   688
TaABI4-1D   GAGACGACTGCCTCGTTGCCAGTGGCTTCGTCATCTACGGTGGCCGCCCAGGATGGTGCGCCGGCGGAGGCGGCAGAGGCTGAGGTGACCCCCGAATGGT   691
Consensus   gagacgactgcc cgtt ccagtggcttcgtcatctacg tggccgcccaggat gtg  ccggcgg ggcggcagaggctgaggtgacccccgaatggt

TaABI4-1A   ACCTTGCCGCCGAGGAGGAGGACTACGAGGCGGCGCTGCTGTGGAATGAACCTGATCCCTTGTTCGACATCTTCTCCAAGTG   782
TaABI4-1B   ACCTTGCCGCCGAGGAGGAGGACTACGAGGCCGCGCTGCTGTGGAATGAACCTGATCCCTTGTTCGACATCTTCTCCAAGTG   770
TaABI4-1D   ACCTTGCCGCCGAGGAGGAGGACTACGAGGCGGCGCTGCTGTGGAATGAACCTGATCCCTTGTTCGACATCTTCTCCAAGTG   773
Consensus   accttgccgccgaggaggaggactacgaggc gcgctgctgtggaatgaacctgatcccttgttcgacatcttctccaagtg
```
