## Supplementary material for "Characterization and expression QTL analysis of *TaABI4*, a pre-harvest sprouting related gene in wheat": TABLE S2

**Table S2.** Primers used for amplification and expressional profile assay of *TaABI4*.

| Name | Sequence (5'-3') | Use of primer |
| --- | --- | --- |
| *TaABI4*-F | CGCCAGGTCCGTCCCTATCT | Forward primer for amplifying *TaABI4* gene |
| *TaABI4*-R | ACCACGACATTACCCCAAAGC | Reverse primer for amplifying *TaABI4* gene |
| *TaABI4*-Q-F | CAGTGGCTTCGTCATCTAC | Forward primer for detecting  expression of *TaABI4* |
| *TaABI4*-Q-R | GATGTCGAACAAGGGATCAG | Reverse primer for detecting  expression of *TaABI4* |
